# Human enhancers harboring specific sequence composition, activity, and genome organization are linked to the immune response

**DOI:** 10.1101/078477

**Authors:** Charles-Henri Lecellier, Wyeth W. Wasserman, Anthony Mathelier

## Abstract

The FANTOM5 consortium recently characterized 65,423 human enhancers from 1,829 cell and tissue samples using the Cap Analysis of Gene Expression technology. We showed that the guanine and cytosine content at enhancer regions distinguishes two classes of enhancers harboring distinct DNA structural properties at flanking regions. A functional analysis of their predicted gene targets highlighted one class of enhancers as significantly enriched for associations with immune response genes. Moreover, these enhancers were specifically enriched for regulatory motifs recognized by TFs involved in immune response. We observed that enhancers enriched for links to immune response genes were more cell type specific, preferentially activated upon bacterial infection, and with specific response activity. Looking at chromatin capture data, we found that the two classes of enhancers were lying in distinct topologically-associated domains and chromatin loops. Our results suggest that specific nucleotide compositions encode for classes of enhancers that are functionally distinct and specifically organized in the human genome.

## 1 Introduction

Gene expression is regulated through many layers, one of which being the regulation of the transcription of DNA segments into RNA. Transcription factors (TFs) are key proteins regulating this process through their specific binding to the DNA at regulatory elements, the TF binding sites (TFBSs) [1]. These regulatory elements are located within larger regulatory regions, the promoters and enhancers [2]. While promoters are situated around transcription start sites (TSSs), enhancers are distal to the genes they regulate. The canonical view is that chromatin conformation places enhancers in close 3D proximity to their target gene promoters through DNA looping [3–5]. High-resolution chromatin conformation capture (Hi-C) technology maps genomic regions in spatial proximity within cell nuclei [6]. The Hi-C technology identified specific genomic neighbourhoods of chromatin interactions, the topologically associating domains (TADs), which represent chromatin compartments that are stable between cell types and conserved across species [7,8].

Studies have shown relationships between the composition of a DNA sequence in guanine (G) and cytosine (C) and chromatin organization, for instance in relation to nucleosome positioning [9,10] and chromatin architecture [11]. DNA sequence composition and other features of promoter regions have been extensively studied, including such key advances as the discovery of CpG islands. The analysis of promoter regions in the human genome was accelerated by the development of the Cap Analysis of Gene Expression (CAGE) technology [12,13], which identifies active TSSs in a high-throughput manner based on 5’ capped RNA isolation. Using CAGE data, a large scale identification of the precise location of TSSs in human [14] led to the classification of promoters into four classes based on G+C content (%GC) [15]. The study highlighted that GC-rich promoters are associated with genes involved in various binding and protein transport activities while GC-poor promoters are associated with genes responsible for environmental defense responses. While promoters overlapping CpG islands are commonly assumed to be ubiquitous drivers of housekeeping genes, comprehensive analysis of CAGE data from > 900 human samples showed that a subset deliver cell type-specific expression [16].

Large-scale computational analyses of enhancer regions have been hampered by a limited set of bona fide enhancers. An advantage of the CAGE technology is its capacity to identify *in vivo*-transcribed enhancers. Specifically, it identifies active enhancer regions in biological samples by capturing bidirectional RNA transcripts at enhancer boundaries [17]. Using this characteristic of CAGE data, the FANTOM5 project identified 65,423 human enhancers, across 1,829 CAGE libraries [16–18]. Sequence property analysis suggested that the enhancers share properties with CpG-poor promoters [17].

As enhancers are distal to the genes they regulate, it is challenging to predict these relationships. Based on cross-tissue correlations between histone modifications at enhancers and CAGE-derived expression at promoters within 1,000 bp, enhancer-promoter links have been shown to be conserved across cell types [19]. As the CAGE technology captures the level of activity for both promoters and enhancers in the same samples, recent studies predicted enhancer targets by correlating the activity levels of these regulatory regions over hundreds of human samples from the FANTOM5 consortium [17, 20]. Predicted enhancer-gene associations were supported by experimental data from ChIA-PET and Hi-C, and eQTL data [17, 20]. Further, [17] unveiled that closely spaced enhancers were linked to genes involved in immune and defense responses. These results stress that predictions of enhancer-promoter associations are critical to decipher the functional roles of enhancers.

Here, we used the G+C content of the sequences of human CAGE-derived enhancer regions to define two classes of enhancers. Based on the enhancer-gene target pairs characterized by both [17] and [20], we showed that the class of enhancers with a lower G+C content was predicted to be functionally associated with genes involved in the immune response. Accordingly, regulatory motifs associated with immune response TFs like NF-*κ*B are enriched in the DNA sequence of the immune response-related set of enhancers. Independent functional analysis of histone modification and CAGE data highlighted a cell type specificity of these enhancers along with their activation upon bacterial infection. Moreover, the class of enhancers enriched for associations with immune system genes was observed with a distinct response activity pattern following cell stimulation in time-course data sets. Finally, we observed that the two classes of enhancers tended to be structurally organized in the human chromosomes within distinct TADs and DNA chromatin loops.

## 2 Materials and Methods

### 2.1 Human enhancers

We retrieved the hg19 positions of the 65,423 human enhancers from phases 1 and 2 of the FANTOM5 project in BED12 format from fantom.gsc.riken.jp/5/datafiles/phase2. 2/extra/Enhancers/human\_permissive\_enhancers\_phase\_1\_and\_2.bed.gz [16–18]. The enhancers were predicted from CAGE experiments performed on 1,829 libraries (http://fantom.gsc.riken.jp/5/datafiles/phase2.2/extra/Enhancers/Human.sample\_name2library\_id.txt). We extracted DNA sequences for regions of 1,001 bp centered at the enhancer mid-points (columns 7–8 of the BED12 file) using the BEDTools [21] and computed the G+C content of the sequences. We considered the distribution of the G+C content of all the enhancers (mean ~ 45%, median ~ 44%, standard deviation ~ 8) to distinguish two classes of enhancers (%GC below the median, class 1; and %GC above the median, class 2; Figure 1a).

**Figure 1:**
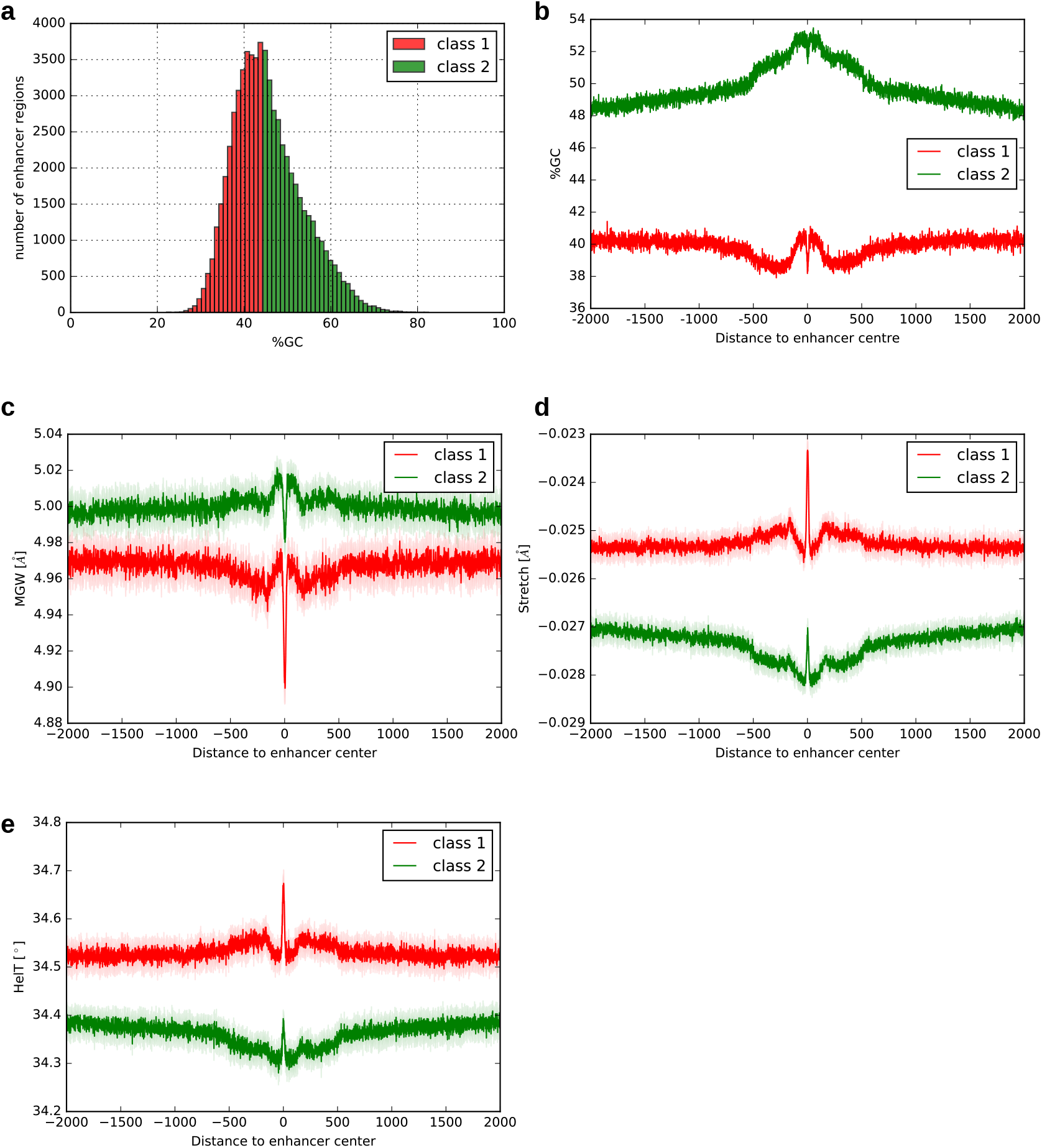
DNA sequence features at enhancers. Features associated with human enhancers from class 1 and class 2 are represented in red and green, respectively. **a**. Histogram of the %GC of the enhancers. **b**. Distribution of the average %GC (y-axis) of the enhancers in classes 1 and 2 along DNA regions of ±2000 bp centered at enhancer center points (x-axis). **c-e**. Average DNA shape values (y-axis) along the DNA regions of ±2, 000 bp centered at enhancer mid-points (x-axis) for DNA shape features MGW (c), Stretch (d), and HelT (e). Shadow regions represent the 90% confidence intervals obtained from bootstrapping (see Materials and Methods). The largest Kolmogorov-Smirnov statistics between class 1 and class 2 enhancers were obtained with these 3 DNA shape features.

### 2.2 Clusterization of human enhancers based on positional distribution of G+C content

Using the DNA sequences for regions of 1,001 bp centered at the enhancer mid-points, we created binary vectors representing the enhancer sequences with 1s and 0s corresponding to G or C and A or T, respectively. Note that 11 enhancers were not considered here as the DNA sequences around their centers contained undefined nucleotides (N in the IUPAC notation). The vectors were clustered using the k-means algorithm implemented in the KMeans function of the *scikit* Python module [52]. The silhouette plots (Figure S1) were constructed for k ∈ [2, 5] using the *silhouette_samples* function of the *scikit* Python module. Formally, the silhouette plots display the silhouette coefficient for each enhancer as (*b* − *a*)/max(*a*, *b*) where *a* is the mean intra-cluster euclidian distance and *b* the mean nearest-cluster euclidian distance.

### 2.3 Distribution of enhancers in the human genome

The distribution of enhancers from the two classes in 3’ UTR, 5’ UTR, intergenic regions, transcription termination sites, intronic regions, non-coding and coding exons, and promoter regions in Figure S3a were obtained using the HOMER (v.4.7.2) *annotatePeaks.pl* script using annotations from the human genome hg19 v.5.4 (http://homer.ucsd.edu/homer/). Distances to TSSs for Figure S3b were obtained using the same script.

### 2.4 Repetitive elements

The hg19 coordinates of repetitive elements were retrieved from the RepeatMasker track of the UCSC Table browser tool (https://genome.ucsc.edu/cgi-bin/hgTables). The overlaps between enhancers and repetitive elements were obtained using the *intersect* subcommand of the BEDTools requiring a minimum overlap of 50% of the enhancer lengths.

### 2.5 Expression quantitative trait loci

The v6p GTEx *cis*-eQTLs (expression quantitative trait loci) were downloaded from the GTEx Portal at http://www.gtexportal.org/home/. The enhancer coordinates from the two classes were intersected with hg19 *cis*-eQTL coordinates using the *intersect* subcommand of the BEDTools. Following *cis*-eQTL variant-gene associations, each enhancer class was linked to potential target genes (2,459 and 5,857 genes for class 1 and class 2 respectively, Table S2). The intersection of the target genes from the two classes yielded 437 genes. To assess the significance of the intersection, we randomly created 1,000 times two classes of enhancers (with 32,487 and 32,936 enhancers, respectively). None of these random selections yielded an intersection of ≤ 437 potential target genes; the corresponding empirical p-value was evaluated as < 10^−3^.

### 2.6 Enhancer gene targets

We considered three sets of enhancer-gene associations. The first two sets were obtained from [17] where enhancer-gene pairs were derived from the correlation of CAGE signal at enhancers and (i) CAGE-derived or (ii) RefSeq-annotated TSSs. The enhancer-CAGE-derived TSSs associations were retrieved from enhancer.binf.ku.dk/presets/human.associations.hdr.txt.gz and the enhancer-RefSeq TSS associations were retrieved from enhancer.binf.ku.dk/presets/ enhancer\_tss\_associations.bed. The third set of enhancer-gene pairs was obtained from [20] where associations were obtained by first computing a lasso-based multiple regression between CAGE signals at each TSS and proximal enhancers from all FANTOM5 samples and then using sample specific information to obtain sample specific pairs. Enhancer-gene pairs were retrieved from http://yiplab.cse.cuhk.edu.hk/jeme/fantom5_lasso.zip. To rely on high quality enhancer-gene pairs, we considered the intersection of the pairs of enhancer-gene targets predicted by the three datasets described above.

We further used the associations from this intersection for promoter sequence analyses. We extracted DNA sequences of ±500 bp around gene start (ENSEMBL hg19 coordinates) using the BEDTools get-fasta [21] and computed average G+C content using the *GC* function of Biopython [22].

### 2.7 MNase profiles

MNase-seq signal from ENCODE for cell lines GM12878 and K562 were obtained as bigWig files from the UCSC genome browser website at http://hgdownload.cse.ucsc.edu/goldenpath/hg19/encodeDCC/wgEncodeSydhNsome/wgEncodeSydhNsomeGm12878Sig.bigWig and http://hgdownload.cse.ucsc.edu/goldenpath/hg19/encodeDCC/wgEncodeSydhNsome/wgEncodeSydhNsomeK562Sig.bigWig. Average MNase-seq signal values at enhancer regions were computed using the *agg* subcommand of the *bwtool* tool [23] with regions spanning ±2,000 bp around the enhancers’ mid-points.

### 2.8 DNA shape feature plots

The values of 13 DNA structural features were retrieved from the GBshape browser [24] as bigwig files at ftp://rohslab.usc.edu/hg19/. We retrieved the averaged DNA shape values at the enhancer regions from class 1 and class 2 using the *agg* sub-command of the *bwtool* tool [23]. The normalized averaged DNA shape values were computed independently for each enhancer class using the equation:

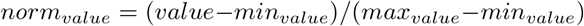

where *norm_value_* is the normalized value to be computed for a DNA shape at a specific position in the DNA sequence, *value* is the averaged DNA shape value at this position for the enhancers in the class, and *min_value_* (*max_value_*) is the minimum (maximum) averaged DNA shape value for the enhancers in the class. The 90% confidence intervals for each DNA shape feature at each position was computed using a bootstrap approach. Specifically, random subsampling of enhancers was used to construct 100 sets of 10,000 randomly selected enhancers from classes 1 and 2. Average DNA shape values were computed for each random set and values from the 5th and 95th percentages were used to define the 90% confidence intervals.

### 2.9 DNA sequence shuffling

The ±2, 000 bp DNA seqnences around enhancer centers were shuffled with the *BiasAway* tool [25]. Specifically, mononucleotide shuffling of the regions was obtained with the *m* subcommand of *BiasAway*.

### 2.10 Gene ontology functional enrichment

To construct Figure 2a, official symbols corresponding to the promoters associated with enhancers from class 1 and class 2 (Table S3) were submitted to GOrilla [26] at http://cbl-gorilla.cs.technion.ac.il/ using the January 7th 2017 update. We used the two unranked lists option with genes associated with enhancers from either class 1, class 2, specific to class 1, specific to class 2, or common to class 1 and class 2 as targets and the aggregated set of genes associated with the full set of enhancers as background. We submitted the enriched GO biological processes with FDR< 0.01 to REVIGO [27] at http://revigo.irb.hr/ asking for a ‘small’ output list. GOrilla outputs are provided in Table S4 and Table S5 for enriched GO biological processes from class 1 associated genes and the intersection of classes 1 and 2 associated genes, respectively.

**Figure 2:**
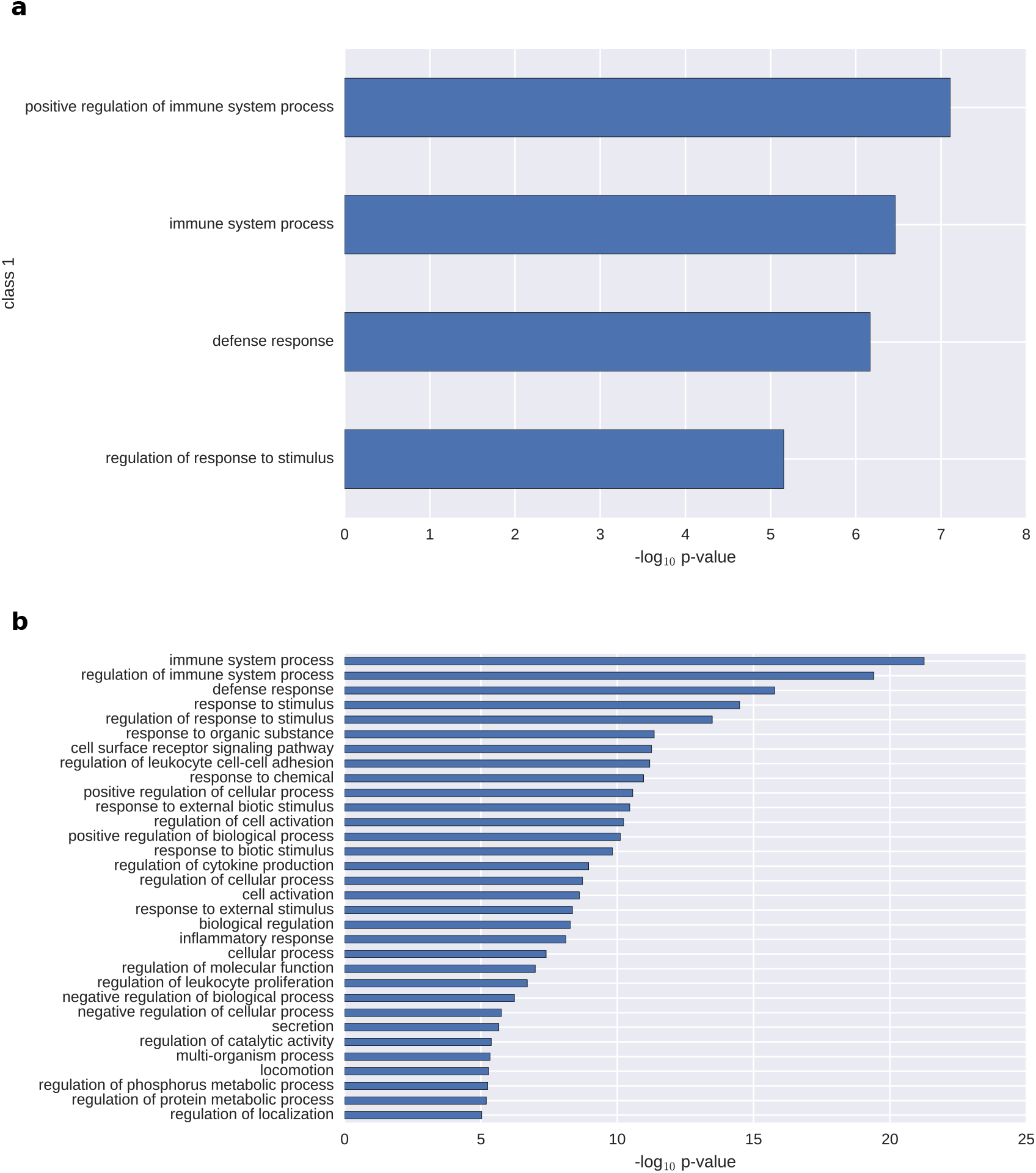
Functional enrichment analysis. Enriched GO biological processes (y-axis; log_10_ p-value < −5) obtained using the GOrilla and REVIGO tools [26, 27] with genes predicted to be regulated by enhancers from class 1 (a) and the ranked list of genes (by decreasing number of associated enhancers) predicted to be regulation by class 1 or 2 enhancers (b). The enhancer-gene pairs were recurrently predicted in three sets of enhancer-gene associations (see Materials and Methods).

To construct Figure 2b, genes were ranked based on the number of enhancers predicted to target them in descreasing order (Tables S6 and S7). This ranked list was submitted to GOrilla using the ranked list option. Enriched GO terms were retrieved as described in the previous paragraph (Table S8).

### 2.11 Motif enrichment

We applied Centrimo [28] from the MEME suite version 4.11.1 with default parameters to DNA sequences of regions ±500 bp around the mid-points of enhancers from class 1 and class 2. Class 1 enhancer regions were used as foreground and class 2 enhancer regions as background and vice-versa. The MEME databases of motifs considered for enrichment were derived from [29] (jolma2013.meme), JASPAR [30] (JASPAR_CORE_2016_vertebrates.meme), Cis-BP [31] (Homo_sapiens.meme), Swiss Regulon [32] (Swiss_Regulon_human_and_mouse.meme), and HOCOMOCO [33] (HOCOMOCOv10_HUMAN_mono meme_format.meme). The same procedure was applied to promoter regions (±500 bp around gene starts) associated with class 1 and class 2 enhancers.

Figure 3a,b has been obtained from the html output of Centrimo by selecting the 3 most enriched motifs (ranked using the Fisher E-value; Data S1–S2, https://doi.org/10.5281/zenodo.1243223).

**Figure 3:**
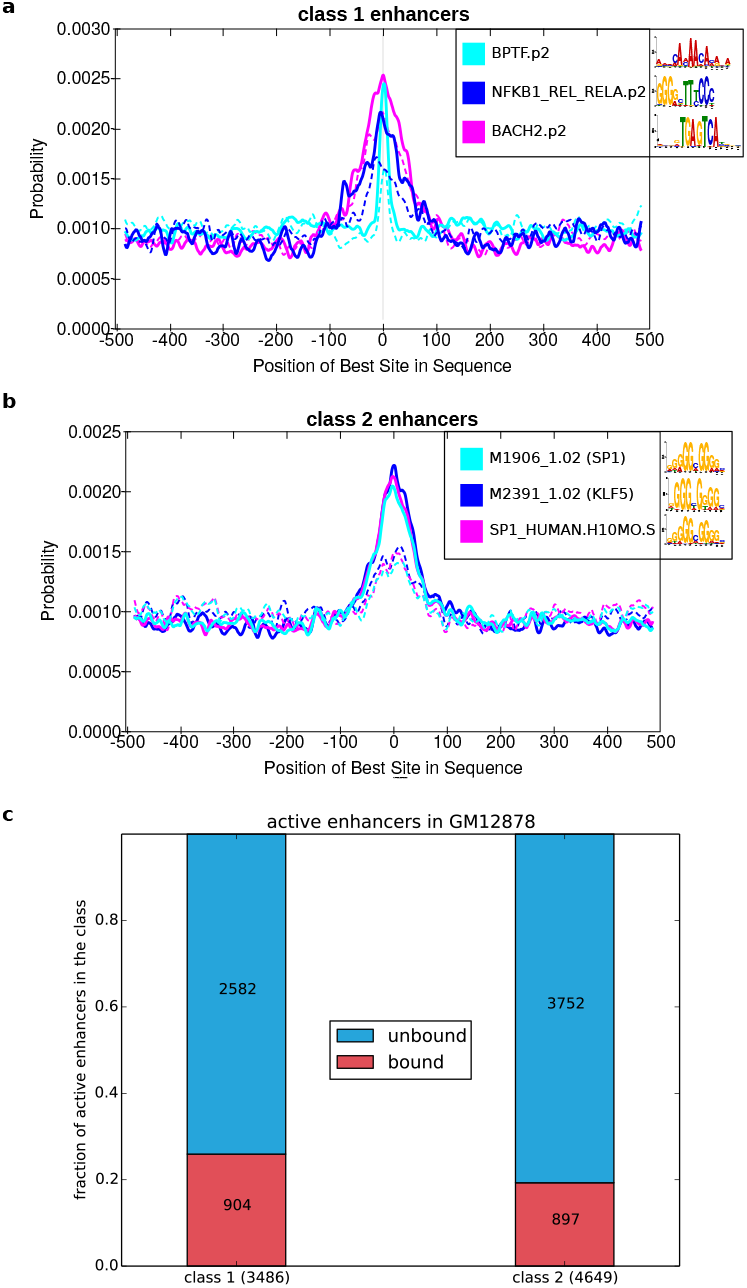
TF binding analysis at enhancer regions. Regions of ±500 bp around enhancer midpoints (**a, b**) were subjected to positional motif enrichment analyses using the Centrimo tool [28] with motifs from JASPAR [30], Cis-BP [31], Swiss Regulon [32], and HOCOMOCO [33]. Enhancers from class 1 (a) and class 2 (b) were analyzed separately. The x-axis represents the distance to the enhancer mid-points. The y-axis represents the probability of predicting TFBSs associated with the motifs given in the legend boxes. Plain lines represent the distribution of predicted TFBSs in the foreground sequences (from class 1 and class 2). Similarly, dashed lines represent the distribution of predicted TFBSs in the background sequences (from class 1 and class 2). Note that the SP1 PWMs enriched in class 2 enhancers originate from [31] (M1906 1.02) and [33] (SP1 HUMAN.H10MO.S). (c) Proportion of class 1 (left) and class 2 (right) active enhancers in GM12878 bound or not by the RELA TF (using ChIP-seq data).

### 2.12 Genome segmentation

#### 2.12.1 ENCODE genome segmentation

The genome segmentation using the combination of results from ChromHMM [34] and Segway [35] for ENCODE tier 1 and tier 2 cell types GM12878, H1hesc, HelaS3, HepG2, HUVEC, and K562 were retrieved from http://hgdownload.cse.ucsc.edu/goldenPath/hg19/encodeDCC/wgEncodeAwgSegmentation/.

#### 2.12.2 Genome segmentation in dendritic cells

The genome segmentation of dentritic cells before and after *Mycobacterium tuberculosis* infection [36] was computed using ChromHMM [34] and retrieved at http://132.219.138.157:8080/DC\_NI\_7\_segments\_modID.bed.gz and http://132.219.138.157:8080/DC\_MTB\_7\_segments\_modID.bed.gz.

#### 2.12.3 Genome segmentation overlap with enhancers

The overlaps between enhancers and genome segments were obtained using the *intersect* subcommand of the BEDTools requiring a minimum overlap of 50% of the enhancer lengths. We considered enhancers as in active states if they overlapped the TSS, promoter flank, enhancer, weak enhancer, and transcribed segments.

### 2.13 RELA ChIP-seq data analyses

The ENCODE RELA ChIP-seq data in GM12878 cells was retrieved at http://hgdownload.cse.ucsc.edu/goldenPath/hg19/encodeDCC/wgEncodeAwgTfbsUniform/wgEncodeAwgTfbsSydhGm12878NfkbTnfaIggrabUniPk.narrowPeak.gz. To identify active FANTOM5 enhancers in GM12878, we considered the overlap between 1,001 bp-long regions around enhancer’s mid-points and genome segments predicted by ChromHMM and Segway combined as enhancer or weak enhancer. The identified 1,001 bp-long active enhancer regions were further overlapped with RELA ChIP-seq peaks. All overlaps were computed with the *intersect* subcommand of the BEDTools.

### 2.14 Enhancer expression specificity

The cell-type expression specificity of enhancers was computed as

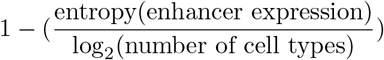

in [17]. Each enhancer expression was represented by a vector of expression values in each cell type, which corresponded to the mean of the enhancer expression in the samples associated with the cell types. The binary matrix of enhancer usage across FANTOM5 samples was obtained at http://enhancer.binf.ku.dk/presets/hg19\_permissive\_enhancer\_usage.csv.gz. The association between FANTOM5 samples and cell types was obtained from Tables S10-S11 in [17]. Heat maps in Figure 6 were computed using the *colormesh* function of the *matplotlib.pyplot* Python module [37].

### 2.15 Enhancer dynamics

FANTOM5 classification in the 14 dynamics displayed in Figure 8 was obtained from Auxiliary data table S3 in [18]. The classification provided response class assignments to 1,294 and 2,800 class 1 and class 2 enhancers, respectively. Response classes were assigned to 2,827 and 4,406 genes associated with class 1 and class 2 enhancers, respectively. Note that enhancers and promoters can be assigned to multiple response classes.

Corresponding plots (Figure 8) and enrichment analyses were performed using *pandas* Python data structure [38] and the *scipy* Python library [39] in the *IPython* environment [40].

### 2.16 Chromatin conformation data

The enrichment for enhancers associated with a specific class in each TAD or chromatin domain (see below) was computed using Binomial test p-values as implemented by the *binom.test* function in the *R* environment [41]. As a control, we randomly assigned the labels class 1 and class 2 to the enhancers and computed the corresponding Binomial test p-values; this procedure was applied to 1,000 random trials. Density plots were obtained using the *density* function in the *R* environment with the *adjust* parameter set to 0.5.

#### 2.16.1 Topologically associating domains

As TADs have been shown to be conserved between cell types and species, we retrieved the TADs defined in the first study describing them [7]. The TADs were predicted in mouse embryonic stem cells and we used the *liftOver* tool from the UCSC genome browser at https://genome.ucsc.edu/cgi-bin/hgLiftOver to map them to hg19 coordinates.

#### 2.16.2 Chromatin loops

The positions of the chromatin loops computed with the HICCUPS tools [42] from Hi-C data on the GM12878, HMEC, HUVEC, HeLa, IMR90, K562, KBM7, and NHEK human cell lines were retrieved from GEO at http://www.ncbi.nlm.nih.gov/geo/query/acc.cgi?acc=GSE63525. Density plots were obtained using the *density* function in the *R* environment with the *adjust* parameter set to 0.5.

### 2.17 Enrichment p-values

P-values throughout the manuscript were computed using the Fisher’s exact test except otherwise stated.

## 3 Results

### 3.1 Guanine and cytosine nucleotide content identified two classes of human enhancers

To analyze the sequence properties of human enhancers, we considered the set of 65,423 CAGE-derived enhancers predicted in the phases 1 and 2 of the FANTOM5 project [16–18]. We extracted 500 bp DNA sequences 5’ and 3’ of the mid-points of the enhancers.

We sought to identify distinct classes of enhancers based on the positional distribution of guanines (Gs) and cytosines (Cs) along the enhancer regions. Specifically, each enhancer was represented by a 1,001 bp-long binary vector with 1s representing G+C and 0s representing adenines (As) and thymines (Ts). We clustered the enhancers by applying the k-means clustering algorithm [43] on the vectors. To select the number of clusters *k*, we considered silhouette plots, which provide a visual representation of how close each enhancer in one cluster is to enhancers in neighbouring clusters [44]. A visual inspection of cluster silhouettes with *k* ∈ [2, 5] revealed that the best clustering was obtained with *k* = 2 (Figure S1). We extracted the two clusters (*k* = 2) of enhancers (containing 42,248 and 23,164 enhancers, respectively) and observed distinct average G+C compositions (Figure S2).

As the clusterization highlighted distinct G+C content between the two classes of enhancers, we next separated the set of enhancers by solely considering their %GC (mean ~ 45%, median ~ 44%, standard deviation ~ 8) to distinguish enhancers with lower (%GC below the median) and higher (%GC above the median) %GC content (Figure 1a). The two classes were composed of 32,487 and 32,936 enhancers, hereafter referred to as class 1 (with lower %GC content) and class 2 (with higher %GC content), respectively. As expected, we observed a large overlap between the two clusters obtained from the k-means algorithm applied to the positional patterns of G+C content along the enhancers and the two classes solely defined from %GC (Jaccard similarity coefficients of 0.77 and 0.7, respectively).

As the mere %GC was sufficient to distinguish classes of enhancers and given the simplicity of this criterion, we used this classification in the following analyses. Next we sought to further explore the positional distribution of the %GC along the enhancer and their flanking regions in classes 1 and 2. We considered ±2, 000 bp DNA sequences 5’ and 3’ of the mid-points of the enhancers and computed the %GC at each position (Figure 1b). We observed distinct positional patterns of G+C content at DNA sequences flanking the enhancers from the two classes. The distinct patterns were emphasized when focusing on the differences in positional patterns of %GC from class 1 and class 2 enhancers represented as the normalized average G+C content separately for the two classes (Figure S5a). This result was expected as the k-means clusterization of the positional patterns of G+C composition provided a classification of enhancers similar to classes 1 and 2. Class 1 enhancers harbored a stronger decrease in %GC at their midpoints when compared to class 2 enhancers. Moreover, the regions surrounding the class 1 enhancers harbored a symmetric decrease in %GC going away from the mid-points with a minimum at about 300–400 bp from the mid-points; it was followed by an increase in %GC. On the contrary, we observed a continuous symmetric decrease in %GC composition going away from class 2 enhancer mid-points. Nevertheless, note that both class 1 and class 2 enhancers harbored a symmetrical decrease of %GC in regions of about 300–400 bp around mid-points.

### 3.2 The two classes of enhancers are associated with distinct interspersed nuclear elements

The two classes of enhancers harbored similar proportions of enhancers located in intronic (~ 55% and ~ 49% of class 1 and class 2 enhancers, respectively) and intergenic (~ 44% and ~ 48% of class 1 and class 2 enhancers, respectively) regions (Figure S3a) but class 2 enhancers were found closer to TSSs than class 1 enhancers (Figure S3b). A third of class 1 enhancers (10,791) and 22% of class 2 enhancers (7,165) overlapped repetitive elements from RepeatMasker. In agreement with their nucleotide composition, class 1 enhancers were enriched in (A)n and (T)n simple repeats and in AT-rich low complexity sequences compared to class 2 enhancers while class 2 enhancers harbored G-rich and C-rich low complexity sequences (Table S1). Further, long interspersed nuclear elements were enriched in class 1 enhancers while no difference was observed for short interspersed nuclear elements (Table S1).

### 3.3 DNA regions flanking the two classes of human enhancers harbored distinct DNA structural properties

As DNA sequence and shape are intrinsically linked, we next considered 13 DNA shape features computed from DNA sequences with the DNAshape tool [45,46]: buckle, helix twist (HelT), minor groove width (MGW), opening, propeller twist (ProT), rise, roll, shear, shift, slide, stagger, stretch, and tilt [47]. We plotted the distribution of these DNA shape features along the enhancers and their flanking regions for the two classes following the same procedure used for analyzing the G+C content (Figures 1ce, S4, and S5). We assessed the pattern differences between class 1 and class 2 enhancers by computing Kolmogorov-Smirnov (K-S) statistics. The three largest K-S statistics were obtained when considering MGW, stretch, and HelT (Figure 1c-e). The main differences between class 1 and class 2 enhancers were observed for regions flanking the enhancers while the regions <~ 200 bp away from the mid-points harbored very similar patterns; this observation was consistent between all 13 DNA shape features (Figures 1c-e and S5b-n). These observations were in agreement with the %GC-patterns observed close to the enhancer mid-points with a symmetric decrease in G+C content and differences when considering flanking regions (Figure 1b). Note that the DNA shape patterns are lost when shuffling the DNA sequences (Figure S6).

Taken together, these results described two subsets of human enhancers distinguishable by their G+C content with distinct positional distribution of %GC along the regions flanking the enhancers, which were reflected in their DNA structural properties. Importantly, we observed that the enhancer classification based on %GC highlighted distinct patterns of DNA shapes along the regions immediately flanking the enhancers but not at the enhancer central regions, indicating that the two classes of enhancers are located in distinct genomic environments.

### 3.4 The two classes of human enhancers associated with distinct biological processes

Different classes of mammalian promoters, derived from their nucleotide composition, were observed to be associated with genes linked to distinct biological functions [15]. Following the same approach, we sought for a functional interpretation of the %GC-based classification of human enhancers. We first aimed at characterizing whether the enhancers from the two classes were associated with distinct sets of target genes based on cis-eQTL associations. We linked enhancers to potential target genes by using cis-eQTL associations defined by the GTEx project. For each enhancer class, a list of potential target genes was obtained for enhancers overlapping with cis-eQTL single nucleotide polymorphisms (see Materials and Methods). We found 2,459 and 5,857 genes linked to class 1 and class 2 enhancers, respectively (Table S2). From these, only 437 genes were linked to enhancers in both classes (p-value < 10^−3^, Fisher’s exact test, see Materials and Methods), suggesting that the %GC-based classification of human enhancers distinguished enhancers regulating different sets of genes.

As the number of enhancer-gene associations is limited from the cis-eQTL analysis (since it requires SNPs in QTL to be located within the CAGE-derived enhancers), we drew enhancer-gene pairs from two previous studies where the pairs were supported by CAGE data using distint approaches [17, 20]. Based on correlations between promoter and enhancer activities derived from CAGE data in human samples, [17] linked enhancers to their potential gene promoter targets. Two sets of associations were computed, based on either CAGE-derived TSSs from [16] or RefSeq-annotated TSSs. More recently, the JEME method [20] used CAGE signal to predict enhancer-gene pairs based on two steps: (1) multiple regression between a TSS and all enhancers in the neighborhood of this TSS across samples, and (2) extraction of sample-specific enhancer-gene pairs. Application of JEME to FANTOM5 samples resulted in a third set of enhancer-gene pairs [20]. Importantly, both FANTOM5 and JEME-based associations have been found to be supported by ChIA-PET, Hi-C, and eQTL data [17,20]. We intersected the predictions of enhancer-gene pairs from these three datasets to rely on high-confidence pairs. We first noticed that G+C content of promoters targeted by class 1 enhancers is lower than that of promoters targeted by class 2 enhancers (Figure S7, Wilcoxon test p-value ¡ 2.2e-16).

To infer the biological functions of enhancers, we assumed that each enhancer was associated with the same biological functions as the genes it was predicted to regulate. We submitted the two sets of genes associated with class 1 and class 2 enhancers to the GOrilla and REViGO tools [26,27] to predict enriched (FDR q-value < 0.01) gene ontology (GO) biological processes. Class 1 enhancers were predicted to target 1,413 genes whereas class 2 enhancers were linked to 2,838 genes (Table S3). In aggregate, the enhancers corresponded to a set of 3,575 genes, of which 676 were common to the two classes (representing ~ 48%, ~ 24%, and ~ 19% of class 1, class 2, and the combined set of genes, respectively). Note that the aggregated set of 3,575 genes was used as the background set of genes for enrichment analyses.

Biological processes linked to immune system processes and response to stimulus were found enriched for genes associated with class 1 enhancers: ‘immune system process’ (q = 6.25 × 10^−9^), ‘positive regulation of immune system process’ (q = 1.68 × 10^−6^), ‘defense response’ (q = 5.49 × 10^−7^, and ‘regulation of response to stimulus’ (q = 3.57 × 10^−4^) (Figure 2a and Table S4). Notably the GO term ‘immune system process’ was found enriched with 647 genes predicted to be targets of class 1 enhancers (Table S4). When considering the genes predicted to be regulated by enhancers from class 2, no GO biological process term was found enriched.

When focusing on the genes predicted to be exclusively targeted by enhancers from class 1 or class 2, we did not find enriched GO terms both for class 1 specific targets and class 2 specific ones. Finally, we considered the set of genes that are predicted to be common targets of enhancers from the two classes. GO terms associated with immune system process, response to stimulus, and cytokine production were found enriched (Table S5). The enrichment of immune system process-related terms was expected as about 48% of the target genes of class 1 enhancers are predicted to be targets of class 2 enhancers as well. It indicates that while immune system genes are targeted by enhancers from both class 1 and class 2, class 1 enhancers are more specifically associated with immune system genes.

Further, we observed that genes linked to the immune response are targeted by a greater number of enhancers than other genes. Indeed, we ranked the list of gene targets of enhancers from classes 1 or 2 by the number of enhancers they were associated with (Table S6) and submitted this list to GOrilla for functional enrichment analysis. This list was derived from 8,339 pairs (Table S7, 2,672 and 5,667 pairs with enhancers from class 1 and class 2, respectively). The GO biological process terms ‘immune system process’, ‘regulation of immune system process’, ‘defense response’, ‘response to stimulus’, and ‘regulation of response to stimulus’ are found at the top of the enriched terms (Figure 2b and Table S8). Furthermore, from the 1,661 enhancer-gene pairs where the gene is associated with ‘immune system process’ (Table S9), 621 were derived from class 1 enhancers, showing that class 1 enhancer-gene associations are enriched in the list of enhancer-gene pairs linked to ‘immune system process’ (Fisher’s exact test p-value = 2.65 × 10^−7^).

Taken together, the functional enrichment results revealed that a classification based on the G+C content of human enhancer regions featured two sets of enhancers predicted to be regulating genes enriched for distinct biological functions. Specifically, while genes linked to ‘immune system process’ were enriched for being linked to a high number of enhancers, the functional enrichments showed that class 1, as opposed to class 2, enhancers were more specifically targeting genes involved in the immune response.

### 3.5 Distinct transcription factors predicted to act upon the two classes of human enhancers

We sought to identify TF binding motifs enriched within each class of enhancers, to suggest driving TFs for the distinct biological functions. We considered 1,001 bp-long DNA sequences centered at the enhancers’ mid-points. Positional motif enrichment analyses were performed using the Centrimo tool [28] to predict TF binding motifs over-represented at enhancers. Class 1 enhancer regions were compared to class 2 regions and vice-versa to highlight specific motifs (Figure 3a,b and Data S1–S2). The most enriched motifs in class 1 enhancer regions were related to the nucleosome-remodeling factor subunit BPTF and the nuclear factor kappa-light-chain-enhancer of activated B cells (NF-*κ*B)/Rel signaling (NFKB1, REL, and RELA [48] and BACH2 [49]; Figure 3a and Data S1–S2), in agreement with an involvement of class 1 enhancers in the immune response biological function (Figure 2). Motifs associated with the Specificity Protein/Krüppel-like Factor (SP/KLF) TFs were enriched in class 2 enhancer regions (Figure 3b and Data S1–S2). Members of the SP/KLF family have been associated with a large range of core cellular processes such as cell growth, proliferation, and differentiation [50]. A similar analysis based on gene promoters associated with class 1 and class 2 enhancers did not yield enriched motifs.

We confirmed the motif-based enrichment of NF-*κ*B/REL/RELA binding in class 1 enhancers by using ENCODE ChIP-seq data obtained in GM12878 cells for the RELA TF involved in NF-*κ*B heterodimer formation. By combining data capturing histone modification marks, TF binding, and open chromatin regions from a specific cell type, the ChromHMM [34] and Segway [35] tools segment the genome into regions associated with specific chromatin states. Focusing on predictions from ChromHMM and Segway combined, we found 3,486 (~ 11%) and 4,649 (~ 14%) active enhancer regions from classes 1 and 2, respectively. We observed that class 1 active enhancers were preferentially bound by RELA. Specifically, 904 active class 1 enhancers and 897 active class 2 enhancers overlapped RELA ChIP-seq peaks (p-value = 1.2×10^−12^, Fisher’s exact test; Figure 3c).

Together, these results reinforced the predictions of biological functions specific to class 1 and class 2 enhancers (Figure 2) through the presence of associated TF binding motifs at enhancers.

### 3.6 The two classes of human enhancers exhibited distinct activity patterns

We further investigated the functional differences between the two classes of human enhancers by analyzing their patterns of activity across cell types. In previous studies, enhancer activity has been inferred either from histone modifications or eRNA transcription signatures [5,34,35,51]. We considered these two approaches. Namely, we considered histone modification data from 6 cell lines and CAGE data from 71 cell types produced by the ENCODE [52] and FANTOM5 [17] projects, respectively.

We retrieved the segmentation of the human genome obtained using a combination of ChromHMM and Segway in the tiers 1 and 2 cell types from ENCODE [52]. For each cell type, we overlapped enhancers with predicted genome segments to assign activity states to the enhancers. As an example, Figure 4 presents the proportion of enhancers from classes 1 and 2 that were overlapping with segments associated with active, CTCF, and repressed chromatin states in embryonic stem cells (H1-hESC). We consistently observed that enhancers from class 2 were significantly more active than those from class 1, which were found to be enriched in repressed genomic segments (Figures 4 and S8). Class 2 enhancers were also associated with segments characterized by CTCF binding.

**Figure 4:**
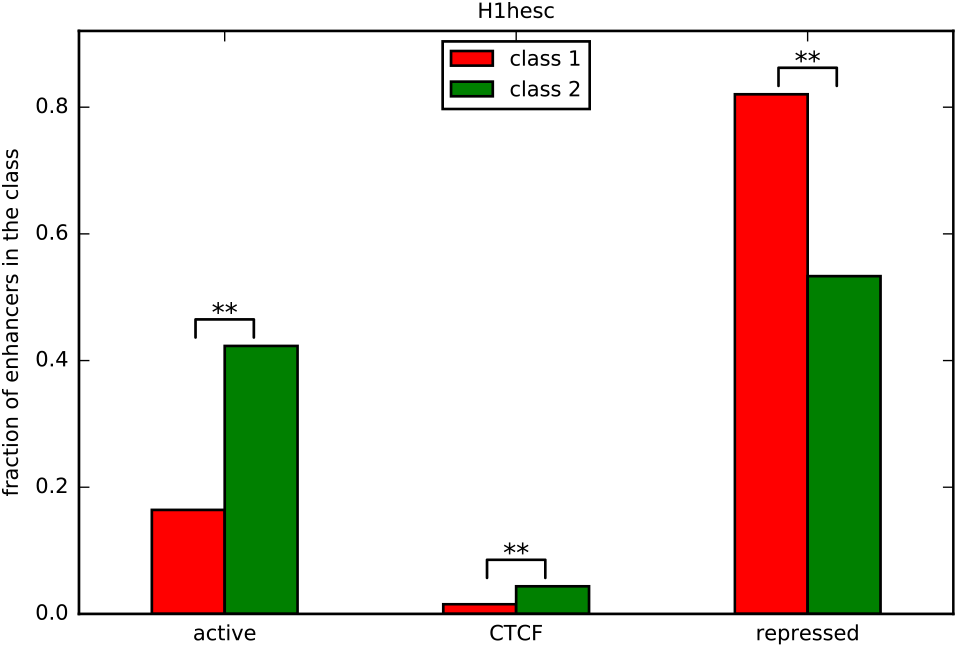
Human enhancers and genome segmentation. Histogram of the proportion of human enhancers (y-axis) in class 1 (red) and class 2 (green) lying within genome segments (x-axis) as annotated by combined predictions from ChromHMM [34] and Segway [35] on human embryonic stem cells (H1-hESC from the ENCODE project [52]). Statistical significance (Bonferroni-corrected p-value < 0.01) of enrichment for enhancers from a specific class is indicated by ‘**’.

To further validate these predictions, we specifically investigated nucleosome occupancy at classes 1 and 2 enhancers and extracted available MNase data at these regions from two cell lines, GM12878 and K562 (Figure 5). Nucleosomes occupancy at class 1 enhancers was found higher than that at class 2 enhancers (Figure 5) in both cell lines, indicating that class 2 enhancers are found more associated with nucleosome depleted regions, in agreement with the fact that they are transcriptionally more active than class 1 enhancers in GM12878 and K562 cells (Figure S8a,e).

**Figure 5:**
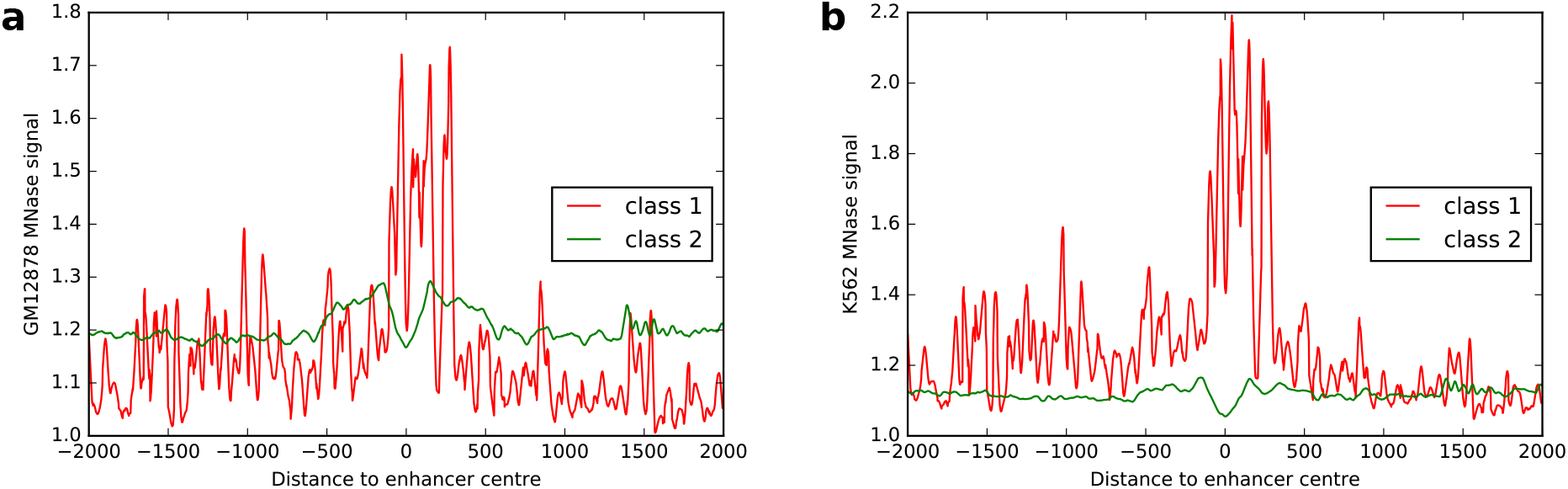
Nucleosome positioning at enhancers. MNase-seq signal from ENCODE for GM12878 (a) and K562 (b) cell lines at regions of ±2, 000 bp around enhancer centers from classes 1 (red) and 2 (green).

From the human samples with CAGE expression from the FANTOM5 project [17], 71 cell types were defined by grouping cell and tissue samples. For each enhancer, [17] computed the entropy of expression of the enhancer across all the cell types. The entropy is then used to compute a cell type-specificity score for each enhancer (see Materials and Methods and [17]). The cell type-specificity score ranges from 0 to 1 with 0 indicating ubiquitous expression and 1 exclusive expression in one cell type. Using this enhancer expression specificity computation, we considered enhancers from class 1 and class 2 separately to highlight potential activity differences in the 71 cell types (Figure 6a-b). Comparing enhancer activity specificity over all the cell types between class 1 and class 2, enhancers from class 1 appeared to be more cell type specific (Figure 6c). While immune cells, neurons, neuronal stem cells, and hepatocytes were previously described to use a higher fraction of human enhancers [17], the elevated utilization was even more pronounced for class 1 enhancers (Figure 6c).

**Figure 6:**
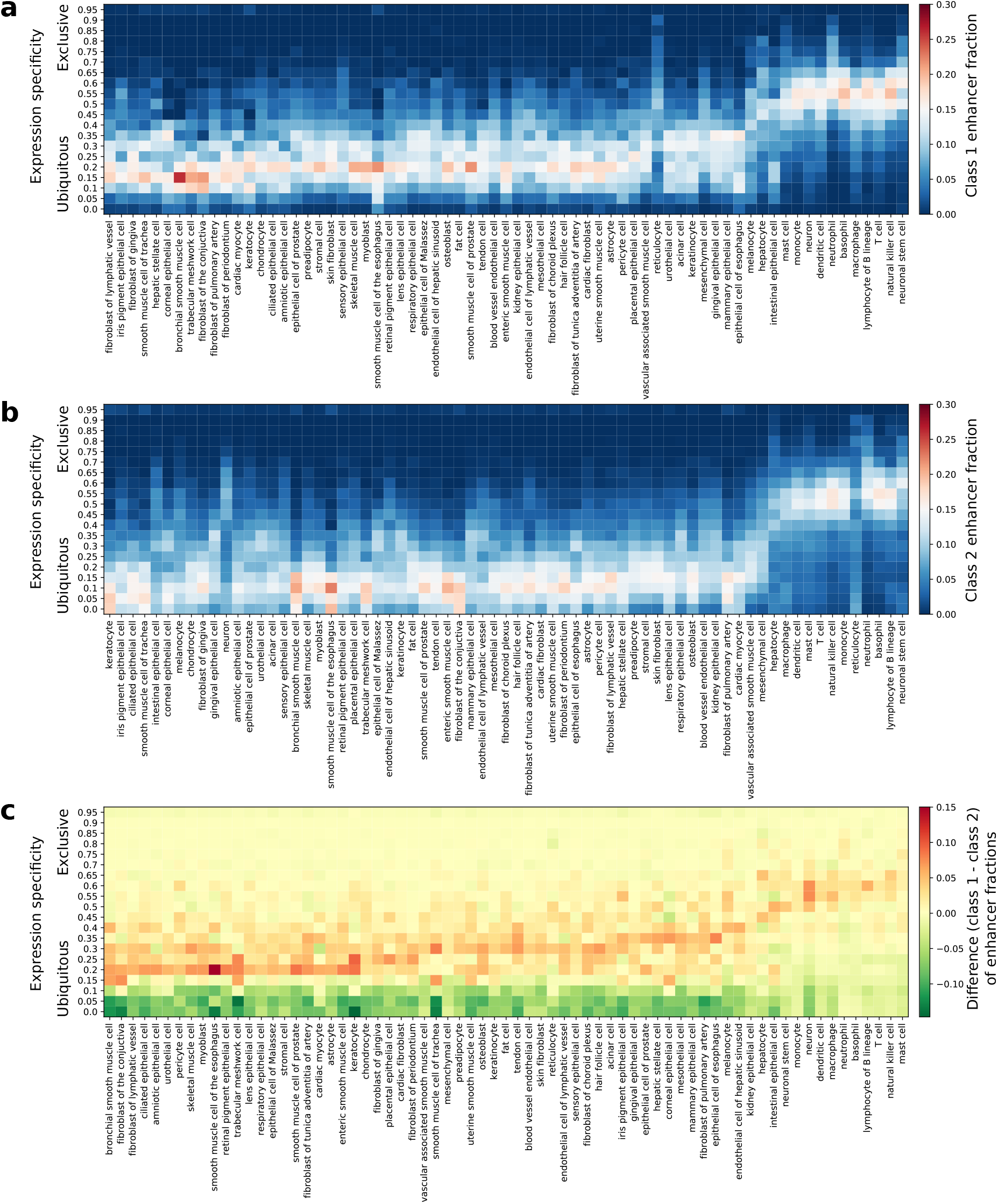
Cell type expression specificities of human enhancers. The cell type expression specificities (see Materials and Methods for details on cell type specificity computation) derived from FANTOM5 CAGE datasets [17] is provided as a heat map for human enhancers in class 1 (**a**) and class 2 (**b**). The colors (see scales) in **a** and **b** represent the fraction of expressed enhancers in each cell type (columns) found in each expression specificity range (rows). The differences in cell type expression specificities between class 1 and class 2 enhancers are provided as a heat map (**c**). Positive (respectively negative) values are represented in red (respectively green) and indicate a higher fraction of class 1 (respectively class 2) enhancers. CAGE, Cap Analysis of Gene Expression.

Taken together, these results derived from histone marks and transcriptional data highlighted that enhancers from class 2 were more ubiquitously active over human cell types than enhancers from class 1, which were more cell type specific. In our previous functional analyses, we inferred the biological functions of the two classes of enhancers from the genes they were predicted to regulate. Here, we further confirmed specific functionalities for the two classes based on enhancer activity analyses, which corroborated with our functional analysis described above. Enhancers from class 1 were more cell type specific, with an emphasis in cell types associated with the immune system, in agreement with the functional enrichment analysis.

### 3.7 Predicted immune system enhancers were activated upon cell infection

We sought to further confirm the association of class 1 enhancers with transcriptional control of immune responses. [36] generated genome-wide DNA methylation, histone marks, and chromatin accessibility data in normal dendritic cells (DCs) and DCs after infection with *Mycobacterium tuberculosis* (MTB). The data provided the opportunity to study the chromatin state changes after infection obtained using the ChromHMM tool [34]. As for the above analysis, we overlapped chromatin state information with the enhancers from classes 1 and 2. To highlight the key epigenetic changes at enhancers, we classified the transition of activities before and after MTB infection into three groups: activated (from inactive before MTB infection to active after infection), inhibited (active to inactive) or unchanged (Figures 7 and S9). We observed that the enhancers from class 1 were significantly more activated (p-value = 4.5 × 10^−8^, Fisher’s exact test) and less inhibited (p-value < 2.2 × 10^−16^, Fisher’s exact test) when compared to class 2 enhancers upon MTB infection (Figure 7). These results reinforced the potential role of class 1 enhancers in immune response.

**Figure 7:**
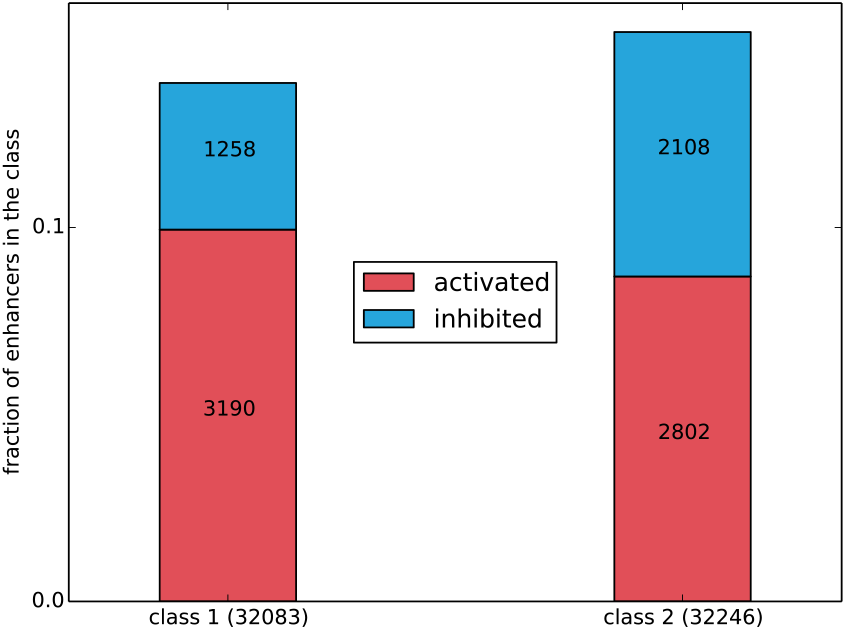
Enhancer activation upon cell infection. Stacked histogram of the fraction of human enhancers (y-axis) from class 1 and class 2 predicted to be activated (red) or inhibited (blue). Predictions were obtained using genomic segments predicted by ChromHMM [34] on human dendritic cells before and after infection with *Myobacterium tuberculosis* [36]. Stacked histogram including unchanged activity is provided in Figure S9.

### 3.8 Predicted immune system enhancers showed specific response activity

Based on time-courses of differentiation and activation, it has been previously reported that transcribed enhancers were coordinating the transcription of genes in transitioning mammalian cells [18]. In this study, CAGE experiments were used to analyze the transcriptional dynamics of enhancers and promoters on the terminal differentiation of stem cells and committed progenitors as well as on the response to stimuli for differentiated primary cells and cell lines [18]. Specifically, they profiled time-courses with CAGE at a high temporal resolution within a 6 hour time-frame to classify enhancers and promoters into distinct dynamic response patterns of early response activity. We overlaid our classification of human enhancers and their predicted target promoters with the response pattern data (Figure 8). Within the enhancers associated with any response pattern (*n* =4, 094; 1,294 and 2,800 from class 1 and class 2, respectively), class 2 enhancers were enriched (p-value = 3.9 × 10^−129^, hypergeometric test).

**Figure 8:**
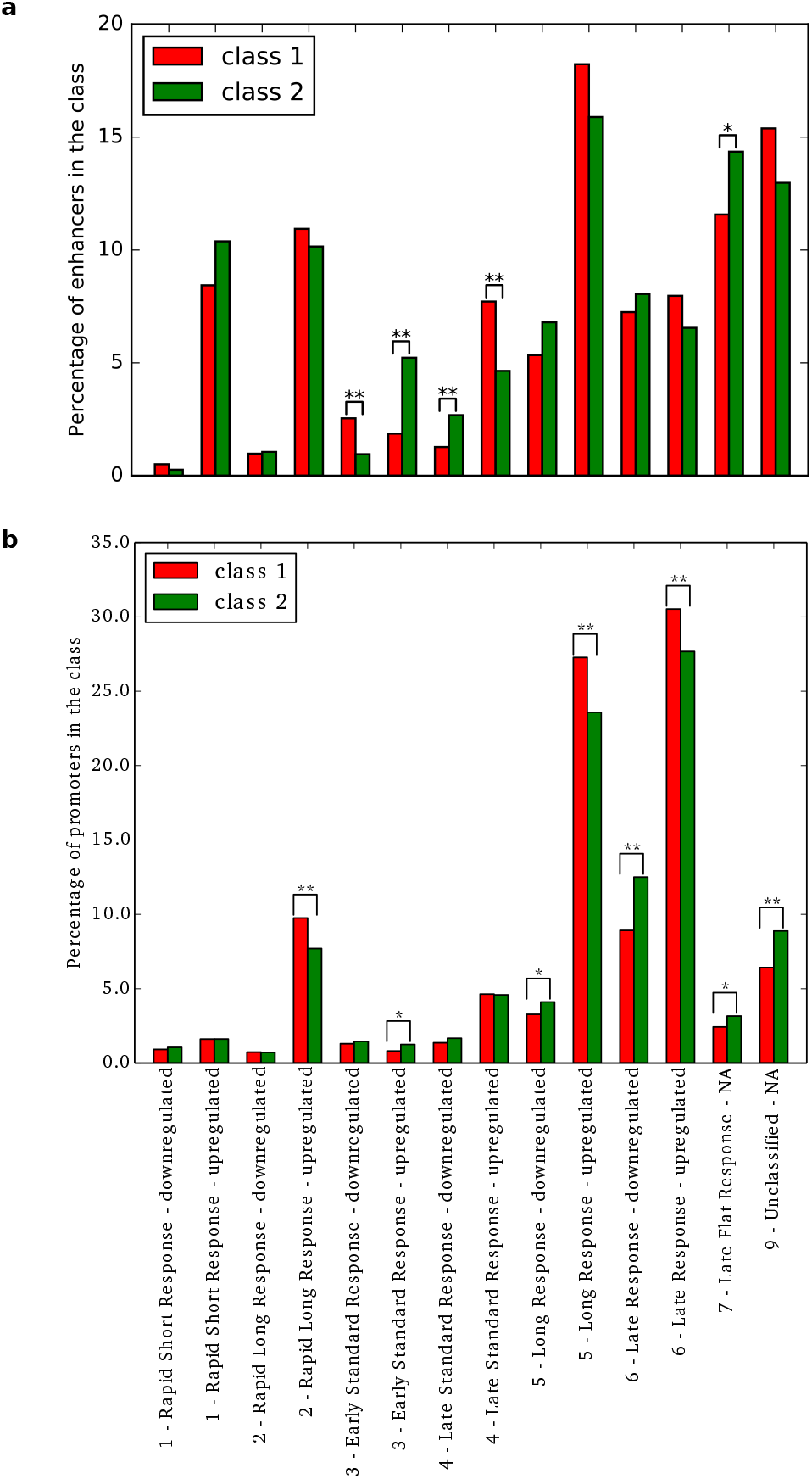
Expression dynamics of human enhancers and associated promoters. Response patterns (x-axis) of human enhancers (**a**) and promoters (**b**) in time courses were classified by [18]. The percentage (y-axis) of enhancers (top) and promoters (bottom) from class 1 (red) and class 2 (green) in each response pattern category are provided as histograms in the two panels. A significant difference between class 1 and class 2 enhancers or promoters in a specific category is highlighted by ‘*’ or ‘**’ for Bonferroni-corrected p-value < 0.05 or < 0.01, respectively (Fisher’s exact tests).

We focused on the set of 4,094 enhancers classified in the dynamic response patterns. Class 1 enhancers were found enriched for down-regulation in ‘early standard response’ and up-regulation in ‘late standard response’. Conversely, class 2 enhancers appeared enriched for up-regulation in ‘early standard response’ and for down-regulation in ‘late standard response’. These results highlighted opposite activity dynamics between class 1 and class 2 enhancers. Note that class 2 was also enriched for up-regulation in ‘late response’. On the other hand, promoters associated to class 1 were enriched in up-regulation of ‘rapid long response’, ‘late response’ and ‘long response’, while promoters associated to class 2 enhancers were enriched in up-regulation of ‘early response’, in down-regulation of ‘late response’ and ‘long response’ and in ‘late flat response’ (Figure 8b), further reinforcing the existence of different transcriptional responses associated with class 1 and class 2 enhancers.

### 3.9 Enhancers from the same class co-localized within chromatin domains

The organization of the chromatin in cell nuclei is a key feature in gene expression regulation by forming regulatory region interactions within TADs [8]. Genes within the same TAD tend to be coordinately expressed across cell types and tissues, and clusters of functionally related genes requiring co-regulation tend to lie within the same TADs [8, 53]. Similar to these studies analyzing gene organization observed in chromatin domains, we focused on how the two classes of enhancers were organized with respect to TADs. We compared the distribution of enhancers from the two classes within a set of TADs [7]. Specifically, we assessed whether individual TADs were biased for containing more enhancers associated with a specific class than expected by chance using the Binomial test. The distribution of the corresponding p-values was compared to those obtained by randomly assigning classes 1 and 2 labels to the enhancers. The results highlighted that TADs were enriched for enhancers from a specific class (Figure 9a), showing a genomic organization of human enhancers with respect to chromatin domains.

**Figure 9:**
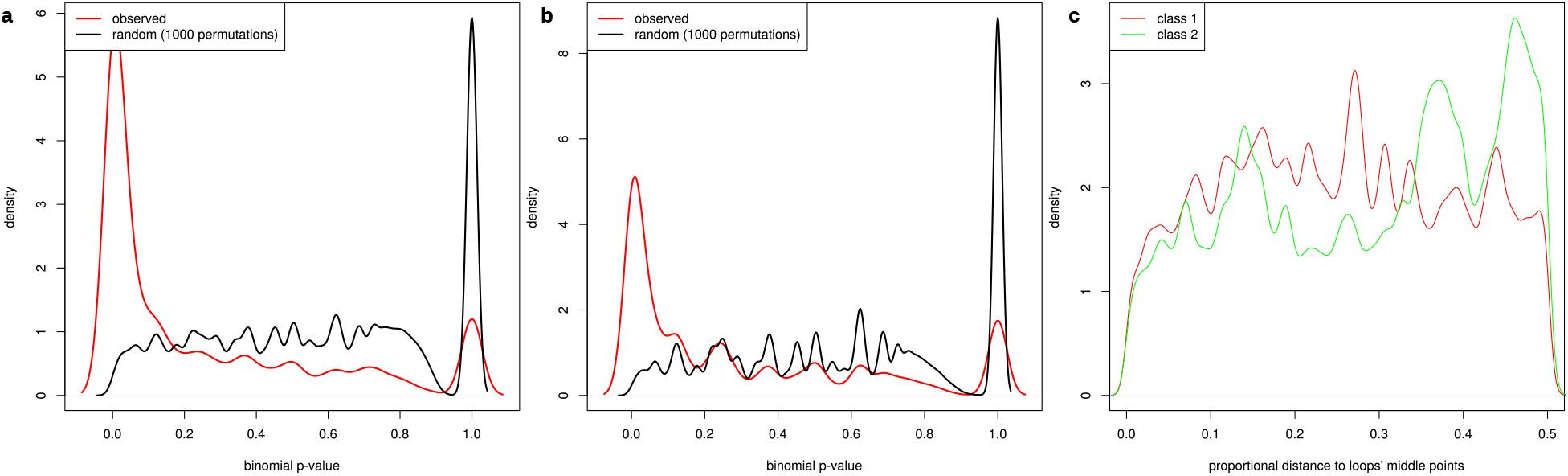
Chromosomal organization of class 1 and class 2 enhancers. **a**. For each TAD [7], we computed the p-value of the Binomial test to assess the enrichment for enhancers from a specific class. The plot compares the density (y-axis) of p-values for Binomial tests (x-axis) applied to classes 1 and 2 enhancers (red) and 1,000 random assignments of class labels to the enhancers (black). **b**. The same analysis as in panel a. was performed using chromatin loops predicted in lymphoblastoid GM12878 cells [42]. **c**. Density (y-axis) of distances (x-axis) between enhancers and chromatin loop centers defined using Hi-C data in GM12878 cells [42]. The distances were normalized by the length of the loops. Enhancers at the center of the loops were found at distance 0.0 while enhancers at chromatin loops boundaries were found at distance 0.5. Results associated with class 1 and class 2 enhancers are depicted in red and green, respectively.

TADs represent interactions within megabase-sized domains of chromatin, which can be subdivided into kilobase-sized chromatin loops of chromatin interactions [42]. We refined our analyses of class-based enhancer co-localization by focusing on chromosomal loops derived from 8 cell lines [42]. Similar to what we observed at the TAD level, we found that chromatin loops tended to contain enhancers from a specific class (Figures 9b and S10). Furthermore, class 1 enhancers were evenly distributed within the chromatin loops whereas enhancers from class 2 were observed to be situated close to the loop boundaries (Figure 9c and S11). This observation is in agreement with the enrichment for class 2 enhancers in CTCF chromatin segments (Figure 4) as chromatin loop boundaries are known to be enriched for CTCF binding [42].

## 4 Discussion

We have analyzed the sequence properties of FANTOM5 human enhancers derived from CAGE experiments to reveal that a subset with lower G+C content is more specifically associated with immune response genes. This set of enhancers tend to co-localize within chromatin domains, exhibit cell type specificity, is activated upon infection, and is observed with specific response activity. In summary, our study of enhancer DNA sequence composition culminated with the identification of human enhancers associated with genes enriched for immune response functions that harbor specific sequence composition, activity, and genome organization.

Note that while immune response genes were found to be targeted by enhancers from both classes, the enhancers with lower %GC were more specifically targeting these genes (Figure 2). As these enhancers are active in a more cell type-specific manner, it suggests that activation of immune response genes in specific cell types is driven by such enhancers bound by NF*κ*B and related TFs (Figure 3).

The analyses of sequence properties in regulatory regions, most prominently CpG islands at promoters, have been key to understanding gene expression regulation [9, 10, 15]. We observed that enhancers with a higher G+C content were more broadly activated than the enhancers with a lower G+C content. A recent study highlighted that human enhancers with broad regulatory activity across cellular contexts were enriched for GC-rich sequence motifs, in line with the fact that broadly active human TFs bind to GC-rich motifs [54]. The enhancers more specifically associated with immune response genes predicted here exhibit a cell-type specific expression pattern and have lower %GC. It remains unclear how and why this set of enhancers has emerged with such sequence properties. In line with their lower G+C content, they were associated with (A)n and (T)n simple repeats and AT-rich low complexity sequences. They were also strongly enriched in long interspersed nuclear elements (LINEs) when compared to other enhancers. Provided that repetitive elements represent both molecular parasites and evolutionary drivers [55], these observations may explain, at the sequence level, the differences observed between the two classes of enhancers. Besides, they suggest that, similar to Alu elements and endogenous retroviruses [56–58], LINEs can exert enhancer activities that are specifically related to the immune response process. In line with this proposal, expression of LINE-1 has been shown to trigger inflammatory pathway in systemic autoimmune disease [59, 60].

When comparing the genomic distribution of the two sets of enhancers, we found that they lie in specific genomic environments associated with a distinct local DNA shape pattern. DNA structural properties were shown to be linked with DNA flexibility, nucleosome positioning, and gene expression regulation [61–65]. We noticed that the DNA shape conformation at enhancers were similar between the two classes but distinct at their flanking regions (Figures 1c-e and S4). The differences in DNA shape features between the two classes of enhancers might relate to differences in conformational flexibility. Indeed, we observed that class 1 enhancer flanking regions harbored increasing MGW, stagger, and opening combined with decreasing HelT close to enhancers compared to class 2 enhancers (Figures 1c-e and S4). These characteristics all relate to distinct flexibility of the DNA, which could provide a topological explanation for the differences observed between the two classes.

One simple explanation for the observation that enhancers from the same class are colocalized would be that enhancers from the same TADs can be found within the same isochore, simply because isochores can help define TADs [11]. For enhancer function, this implies that enhancer-promoter associations can be governed by sequence-level instructions, like G+C content. This idea is in line with the existence of a sequence-encoded enhancer-promoter specificity as unveiled by [66] and [67], which is also supported by our findings related to the association of enhancers with gene promoters of similar %GC composition (Figure S7).

## 5 Acknowledgments

As research parasites [68], we are indebted to the researchers around the globe who generated experimental data and made them freely available. We thank Miroslav Hatas and Georgios Magklaras for systems support, Dora Pak for management support, Chih-Yu Chen for providing the code for enrichment analyses in chromatin conformation data, Tsu-Pei Chiu, Jinsen Li, and Remo Rohs for providing DNA shape feature computation before their publication and their help in interpreting the DNA shape results, Robin Andersson for his help with FANTOM5 enhancer data, and Vladimir Teif for insightful discussion on nucleosome positioning. CHL was supported by funding from CNRS, *Plan d’Investissement d’Avenir* #ANR-11-BINF-0002 *Institut de Biologie Computationnelle* (Young Investigator grant). AM and WWW were supported by the Genome Canada/Genome BC Large Scale Applied Research Grant 174CDE. Funding was provided by the Child and Family Research Institute and the British Columbia Children’s Hospital Foundation, Vancouver, to AM and WWW. AM was also supported by funding from the Norwegian Research Council (Project Number 187615), Helse Sør-Øst, and the University of Oslo through the Centre for Molecular Medicine Norway (NCMM) and the Oslo University Hospital Radiumhospitalet.

## 6 Author contributions

CHL and AM conceived and designed the project. CHL and AM implemented and performed experiments. CHL, WWW, and AM analyzed and interpreted the results. CHL and AM wrote the manuscript with revisions from WWW.

## 7 Supplementary Figure Legends

Figure S1: Silhouette plots of k-means clusters. Silhouette plots for clusters obtained using the k-means clusterization algorithm for *k* = 2 (a), *k* =3 (b), *k* = 4 (c), and *k* = 5 (d). After clusterization of the enhancers using the k-means algorithm, the silhouette score was computed for each enhancer vector. For each k-means clusterization corresponding to each panel, clusters are represented with different colors. A silhouette score can range from −1 to 1 with −1 indicating a possible assignment of the enhancer vector to the wrong cluster, 0 indicating that the vector is close to the boundary between two clusters, and 1 indicating that the vector is far away from the boundary between two clusters. The silhouette score is calculated as (*b* − *a*)*/*max(*a*, *b*) where *a* is the mean intra-cluster distance and *b* the mean nearest-cluster distance for each sample as implemented in the scikit-learn silhouette score function. The red dashed lines represent the average silhouette score over all the enhancer vectors.

Figure S2: %GC composition of enhancers in k-means clusters. Histogram of the %GC of the enhancers in the two clusters (represented with red and green colors) obtained from the k-means algorithm applied to the positional patterns of G+C along human enhancers.

Figure S3: Localization of the enhancers from the two classes. (a) Proportion of enhancers from class 1 (left) and 2 (right) located in 3’ UTR, 5’ UTR, intergenic regions, transcription termination sites (TTS), intronic regions, non-coding and conding exons, and promoter regions. (b) Density distribution of distances to transcription start sites (TSSs) for enhancers from class 1 (red) and class 2 (green).

Figure S4: DNA shape patterns at enhancers (to be continued). Features associated with human enhancers from class 1 and class 2 are represented in red and green, respectively. **a-j** Average DNA shape values (y-axis; full lines) along the ±2, 000 bp DNA sequences centered at enhancer mid-points (x-axis) for DNA shape features buckle (a), opening (b), ProT (c), rise (d), roll (e), shear (f), shift (g), slide (h), stagger (i), and tilt (j). Shadowed areas represent the 90% bootstrap confidence intervals.

Figure S5: Normalized %GC and DNA shape patterns at enhancers (to be continued). Features associated with human enhancers from class 1 and class 2 are represented in red and green, respectively. Distribution of the normalized average %GC (a) and DNA shape values (y-axis) along the ±2, 000 bp DNA sequences centered at enhancer midpoints (x-axis). DNA shapes considered are buckle (b), HelT (c), MGW (d), opening (e), ProT (f), rise (g), roll (h), shear (i), shift (j), slide (k), stagger (l), stretch (m), and tilt (n).

Figure S6: DNA shapes from shuffled enhancer sequences. DNA shape values for HelT (a), MGW (b), ProT (c), and Roll (d) computed using the DNAshapeR tool from shuffled sequences around class 1 (red) and class 2 (green) enhancers.

Figure S7: G+C content of promoters associated to each class of enhancers. We extracted sequences spanning 500 bp around the start of genes associated to class 1 and class 2 enhancers (based on FANTOM5 + JEME associations) and computed GC content using the *GC* function of Biopython (median = 59.04 and 62.14 for promoters associated to respectively class 1 and class 2 enhancers, Wilcoxon test p-value < 2.2e-16).

Figure S8: Human enhancers and genome segmentation. Histograms of the proportion of human enhancers (y-axis) in class 1 (red) and class 2 (green) lying within genome segments (x-axis) as annotated by combined results of ChromHMM and Segway on human lymphoblastoid (GM12878; **a**), cervical cancer (HeLa-S3; **b**), liver carcinoma (HepG2; **c**), umbilical vein endothelial (HUVEC; **d**), and chronic myelogenous leukemia (K562; **e**) cell lines from the ENCODE project. Statistical significance (Bonferroni-corrected Fisher exact test p-value < 0.01) of enrichment for enhancers from a specific class is indicated by ‘**’.

Figure S9: Enhancer activation upon MTB infection. Stacked histogram of the fraction of human enhancers (y-axis) from class 1 and class 2 predicted to be activated (red), inhibited (blue), or with unchanged activity (grey). Predictions were obtained using genomic segments predicted by ChromHMM on human dendritic cells before and after infection with Myobacterium tuberculosis.

Figure S10: Enrichment of enhancers from a single class within chromatin loops. For each chromatin loop predicted in HeLa (**a**), HMEC (**b**), HUVEC (**c**), IMR90 (**d**), K562 (e), KBM7 (**f**), and NHEK (**g**) cell lines, we computed the p-value of the Binomial test to assess enrichment for enhancers from a single class. The plots compare the density (y-axis) of p-values for Binomial tests (x-axis) applied to class 1 and 2 enhancers (red) and 1,000 random assignments of class labels to the enhancers (black).

Figure S11: Organization of enhancers within chromatin loops. Density (y-axis) of distances (x-axis) between enhancers and chromatin loop centers/anchors. The distances were normalized by the lengths of the loops. Enhancers at the center of the loops were found at distance 0.0 while enhancers at chromatin loops boundaries/anchors were found at distance 0.5. Results associated with class 1 and class 2 enhancers are depicted in red and green, respectively. HeLa (**a**), HMEC (**b**), HUVEC (**c**), IMR90 (**d**), K562 (**e**), KBM7 (**f**), and NHEK (**g**) cell lines were considered. In all cell types tested except Hela, class 1 enhancers were evenly distributed within the chromatin loops whereas enhancers from class 2 were observed to be situated close to the loop boundaries.

